# Dynamic Control of Prokaryotic Chromosome Ploidy Rewires Metabolic Networks to Enhance Product Biosynthesis

**DOI:** 10.64898/2026.08.27.747416

**Authors:** Xin Jin, Yaping Gao, Hannuo Shen, Xuanye Zhang, Xiaomei Xu, Sumeng Wang, Qingsheng Qi, Quanfeng Liang

## Abstract

Building high-performance microbial cell factories requires dynamic coordination of resource allocation among cellular growth, target-product biosynthesis, and endogenous host metabolism. However, existing polyploid engineering strategies rely primarily on static manipulation of chromosome copy number. Although increasing gene dosage can enhance biosynthetic capacity, static designs cannot readily accommodate the changing metabolic demands encountered during fermentation. Here, we developed a metabolite-responsive dynamic polyploid engineering strategy that couples chromosome ploidy to the cellular metabolic state. We first constructed a high-performance L-threonine biosensor and used it to sense intracellular L-threonine levels and regulate *ftsZ* expression, a key cell-division gene, thereby establishing a dynamic polyploid system that requires neither exogenous inducers nor antibiotics. This system enabled engineered cells to progressively transition from polyploid to haploid during fermentation, accompanied by stage-specific remodeling of cellular physiology and metabolism. Physiological characterization revealed a marked increase in cell size and alterations in cell-envelope properties during the polyploid phase, followed by a gradual decrease in chromosome copy number as fermentation progressed. Transcriptomic and metabolomic analyses further demonstrated that dynamic ploidy transitions induced global metabolic network rewiring, remodeling the tricarboxylic acid cycle and amino acid metabolism while redirecting carbon flux toward the biosynthesis of aspartate-family amino acids. Ultimately, dynamic polyploid engineering substantially enhanced L-threonine production, enabling the engineered strain to achieve an L-threonine titer of 183.1 g/L and a yield of 0.67 g/g glucose in 5-L fed-batch fermentation without antibiotics or exogenous inducers. These findings show that dynamic regulation of chromosome ploidy can couple gene-dosage control with remodeling of cellular physiology and metabolic networks, providing a new engineering strategy to overcome the limitations of static polyploid designs and build high-performance microbial cell factories.

## Introduction

Industrial biomanufacturing harnesses microorganisms to convert renewable carbon feedstocks into value-added chemicals, offering an important route to reduce dependence on fossil resources and advance sustainable manufacturing^1–3^. A central challenge in building high-performance microbial cell factories is allocating cellular resources efficiently among growth, host homeostasis, and target-product biosynthesis^4^. Conventional metabolic engineering typically enhances pathway flux by increasing the copy number of genes encoding key enzymes, strengthening promoter activity, or attenuating competing pathways^5^. However, sustained high-level gene expression competes for limited transcriptional, translational, and energetic resources, potentially imposing a substantial metabolic burden, impairing growth, and compromising genetic stability^6–8^. More importantly, microbial fermentation is inherently dynamic: cellular growth states, substrate availability, and metabolite levels change continuously throughout the process, whereas static engineering designs cannot readily align engineered gene expression with stage-specific metabolic demands. Coordinating resource allocation between biomass formation and product synthesis over time therefore remains a major challenge in improving the performance of microbial cell factories^9, 10^.

To overcome the limitations of static metabolic engineering^11, 12^, researchers have increasingly used dynamic metabolic control to modulate key pathways expression in response to cellular states or metabolite levels, thereby reallocating metabolic resources across growth phases^13, 14^. Nevertheless, conventional flux optimization and biosensor-and regulatory circuit-based dynamic control have focused primarily on enzyme activity, gene expression, or individual metabolic pathways^15^. By contrast, chromosome copy number determines genome-wide gene dosage. It is closely associated with transcriptional and translational capacity, cellular physiology, and metabolic network organization^16, 17^, potentially representing an additional engineering layer distinct from local flux modulation and transcriptional regulation. From this perspective, manipulating chromosome ploidy to alter a cell’s genomic state could coordinately modulate multiple gene dosages and reshape host metabolism at the chromosome scale, introducing a new dimension for controlling resource competition within complex metabolic networks.

Polyploid engineering provides a foundation for exploring this concept. Increasing chromosome copy number simultaneously elevates genome-wide gene dosage and alters cell size, transcriptional and translational activity, and metabolic state, conferring a form of global regulation fundamentally distinct from single-gene overexpression or local pathway optimization^17–19^. However, existing engineered polyploid systems generally employ static designs with fixed chromosome copy numbers. Consequently, both the enhanced biosynthetic potential conferred by elevated gene dosage and the resource expenditure required to maintain multiple genome copies persist throughout fermentation. Chromosome replication demands a continuous supply of nucleotides, energy, and replication-associated machinery^20, 21^. Thus, as cellular metabolic demands shift during the later stages of fermentation, maintaining high ploidy may no longer provide an advantage in resource allocation. This raises an unresolved question: can chromosome ploidy be transformed from a fixed cellular property into a dynamically tunable engineering variable, allowing gene dosage to change in accordance with fermentation stage and metabolic state?

To address this question, we developed a metabolic-state-driven strategy to regulate chromosome ploidy dynamically. By temporally modulating chromosome copy number, this strategy enables cells to maintain a polyploid state during the early stage of fermentation, thereby increasing gene dosage and expanding biosynthetic capacity. As the target product accumulates, chromosome copy number is progressively reduced, avoiding the need to sustain high ploidy throughout the entire fermentation process. This dynamic “polyploid-first, reduced-ploidy-later” regime effectively realigns gene dosage with changing cellular metabolic demands over time, transforming chromosome ploidy from a static genetic attribute into a dynamic engineering parameter that actively participates in fermentation control. Coupling metabolic state to ploidy transitions requires a regulatory module that converts intracellular metabolite concentrations into signals that control cell division. We therefore employed a metabolite-responsive biosensor to link target-product accumulation to chromosome-ploidy transitions, eliminating dependence on exogenous chemical inducers and improving compatibility with industrial fermentation processes. In this study, we established an intracellular-metabolic-state-driven dynamic polyploid engineering system in an industrial L-threonine-producing *Escherichia coli* strain. We first constructed and optimized an L-threonine-responsive biosensor and used it to dynamically regulate *ftsZ*, a key cell-division gene dynamically. This design coupled L-threonine accumulation to chromosome-ploidy transitions, enabling the engineered strain to shift progressively from polyploid to haploid during fermentation. Physiological characterization further revealed changes in cell morphology and physiological state associated with the ploidy transition. Transcriptomic and metabolomic analyses indicated that dynamic ploidy regulation was associated with global metabolic network rewiring and redirected carbon flux toward L-threonine biosynthesis. Ultimately, the engineered strain achieved an L-threonine titer of 183.1 g/L and a sugar-to-product yield of 0.67 g/g in 5-L fed-batch fermentation without antibiotics or exogenous inducers. Collectively, these results demonstrate that chromosome ploidy can be transformed from a conventionally static cellular attribute into a dynamically adjustable engineering variable responsive to metabolic state, providing a new engineering dimension for coordinating gene dosage, cellular physiology, and metabolic networks at the chromosome scale.

## Results

### Construction and validation of a dynamic chromosome-ploidy control system in *Escherichia coli* MG1655

To determine whether chromosome ploidy could serve as a dynamically tunable engineering variable, we first established an inducible ploidy-control system in the model strain *Escherichia coli* MG1655 by regulating *ftsZ*, a core cell-division gene (Fig. 1a). Expression modules containing a chloramphenicol-resistance marker and transcriptional and translational regulatory elements of different strengths were integrated immediately upstream of the *ftsZ* start codon, placing *ftsZ* expression on the engineered chromosome under IPTG-responsive control (Fig. 1b). Under chloramphenicol selection in the absence of IPTG, the engineered chromosome was retained through its resistance marker; however, its low level of *ftsZ* expression was insufficient to support normal cell division independently. Cells were therefore required to retain a wild-type chromosome copy capable of expressing *ftsZ* at a sufficient level, resulting in a multicopy state comprising both engineered and wild-type chromosomes. Following IPTG addition, *ftsZ* expression from the engineered chromosome increased sufficiently to support normal Z-ring formation and cell division. This relieved the selective constraint favoring maintenance of the multicopy state, allowing chromosome content to decrease progressively toward the near-haploid control level. We thus established an artificial control system in which modulation of *ftsZ* expression drives chromosome-ploidy transitions.

**Fig. 1.**
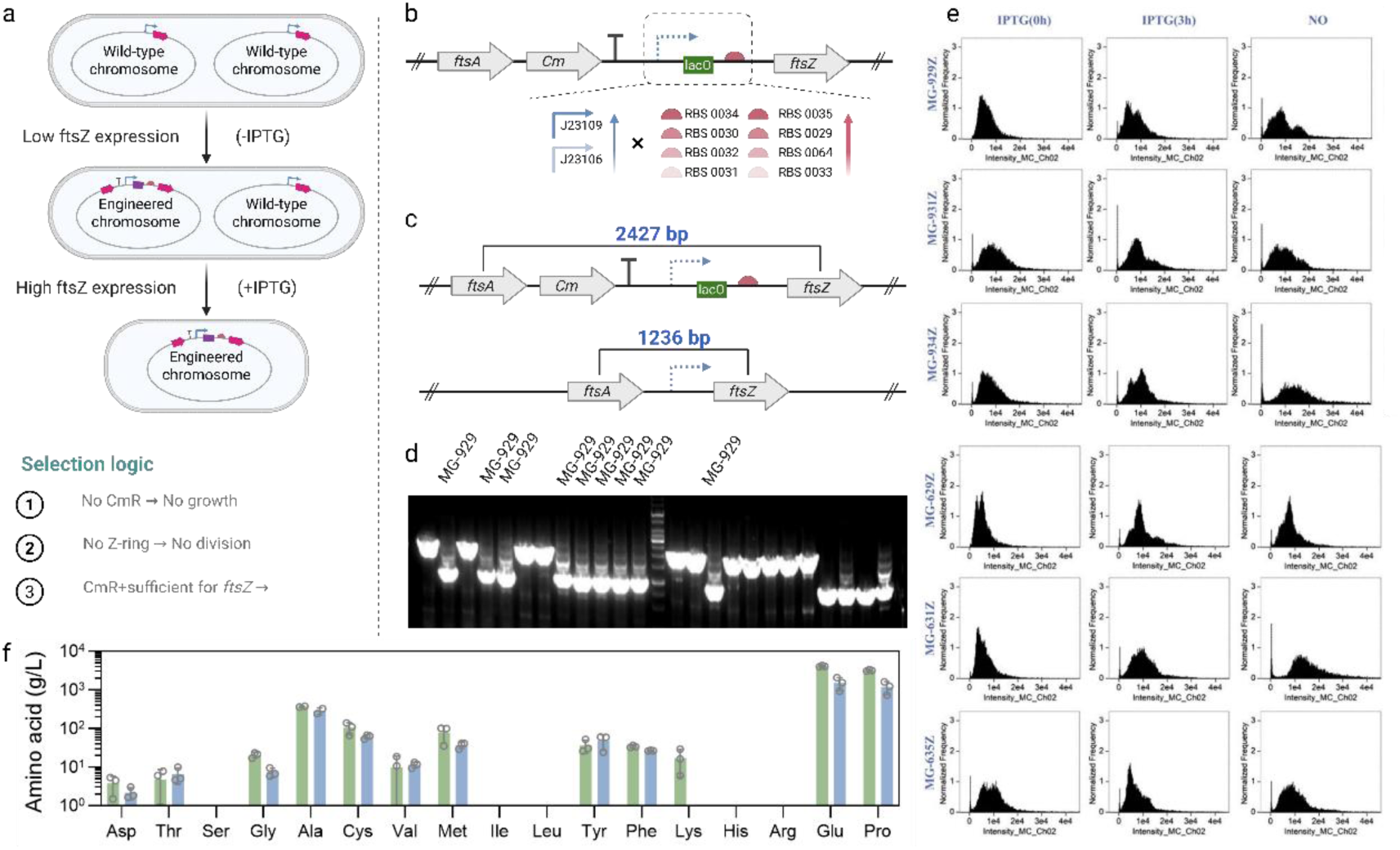
Construction of a dynamic polyploidy system in the model strain *E. coli* MG1655. a, Principle of dynamic chromosome-ploidy regulation. b, Schematic of the genetic construction. c, Validation of the genotype and recombination locus. d, Agarose gel electrophoresis of PCR products. e, Flow-cytometric analysis of cellular DNA content. f, Amino acid-production profiles of MG-631Z and MG1655.

To determine whether this design generated controllable changes in chromosome ploidy, we first characterized the engineered *ftsZ* locus. PCR amplification across the recombination site detected characteristic products corresponding to both the engineered and wild-type *ftsZ* loci in MG-929Z, MG-931Z, MG-934Z, MG-629Z, MG-631Z, and MG-635Z. These results were consistent with the coexistence of the two chromosome types within the respective cell populations (Fig. 1c, d). We subsequently quantified cellular DNA content by fluorescent nucleic-acid staining followed by flow cytometry. Relative to MG1655, MG-929Z, MG-934Z, MG-629Z and MG-631Z showed pronounced population shifts toward higher DNA content in the absence of IPTG, with fluorescence signals approximately two-to threefold higher than those of MG1655. After 3 h of IPTG induction, the flow-cytometric peaks shifted markedly toward lower fluorescence intensities and approached the DNA-content distribution of MG1655 (Fig. 1e). These results demonstrate that modulation of *ftsZ* expression can drive a dynamic transition from a high-chromosome-content state to a low-chromosome-content state in *E. coli*, supporting the feasibility of engineering chromosome ploidy as a dynamically controllable cellular property.

We next investigated whether chromosome-ploidy transitions could alter the host metabolic phenotype. We cultivated MG-631Z, which showed the most pronounced ploidy transition, in M9 medium for 48 h and then quantified its amino acid production profile. Compared with MG1655, MG-631Z exhibited distinct changes in the accumulation of individual amino acids. The concentration of L-threonine increased from 4.9 to 6.6 mg/L, representing an increase of 34.69%; L-valine increased from 9.8 to 11.5 mg/L, an increase of 17.34%; and L-tyrosine increased from 36.4 to 47.9 mg/L, an increase of 31.59%. By contrast, glycine, L-glutamate, and L-proline decreased by 64.73%, 63.27%, and 62.65%, respectively. In contrast, L-lysine fell below the limit of detection (Fig. 1f). These divergent responses indicate that altered chromosome ploidy does not simply enhance overall metabolic activity but instead selectively influences the biosynthesis and accumulation of different amino acids, suggesting broad remodeling of the cellular metabolic state.

Collectively, the dynamic ploidy-control system established in MG1655 enabled a controllable transition from a high-chromosome-content state to a low-chromosome-content state and showed that chromosome-ploidy changes can markedly alter the metabolic output of the host. These findings indicate that chromosome ploidy is not merely a genetic characteristic of the cell but can also serve as an engineerable layer of global regulation, influencing metabolic phenotypes through chromosome-scale changes in gene dosage and cellular resource allocation. This model system therefore provides proof of concept for converting externally induced ploidy control into an autonomous regulatory system responsive to intracellular metabolic states and for applying dynamic chromosome-ploidy engineering to industrial production strains.

### Construction and validation of a metabolite-responsive dynamic chromosome-ploidy control system in the L-threonine-producing strain TG

In the model strain MG1655, we demonstrated that dynamic modulation of *ftsZ*, a key cell-division gene, could drive chromosome-ploidy transitions. However, this system relied on exogenous IPTG induction, limiting its applicability in production strains and large-scale fermentation^22^. We therefore transferred this ploidy-control strategy to the L-threonine-producing strain TG. We used intracellular L-threonine as an endogenous regulatory signal, thereby coupling chromosome-ploidy transitions to target-product biosynthesis. Specifically, the L-threonine-responsive regulatory element ThrSen1.0 was used to control *ftsZ* expression and was combined with ribosome-binding sites (RBSs) of different translational strengths to generate candidate strains with distinct ploidy-transition kinetics (Fig. 2a). This design replaced exogenous chemical induction with dynamic control driven by intracellular metabolite accumulation, providing a basis for adjusting chromosome ploidy in accordance with the metabolic state of the production strain.

**Fig. 2.**
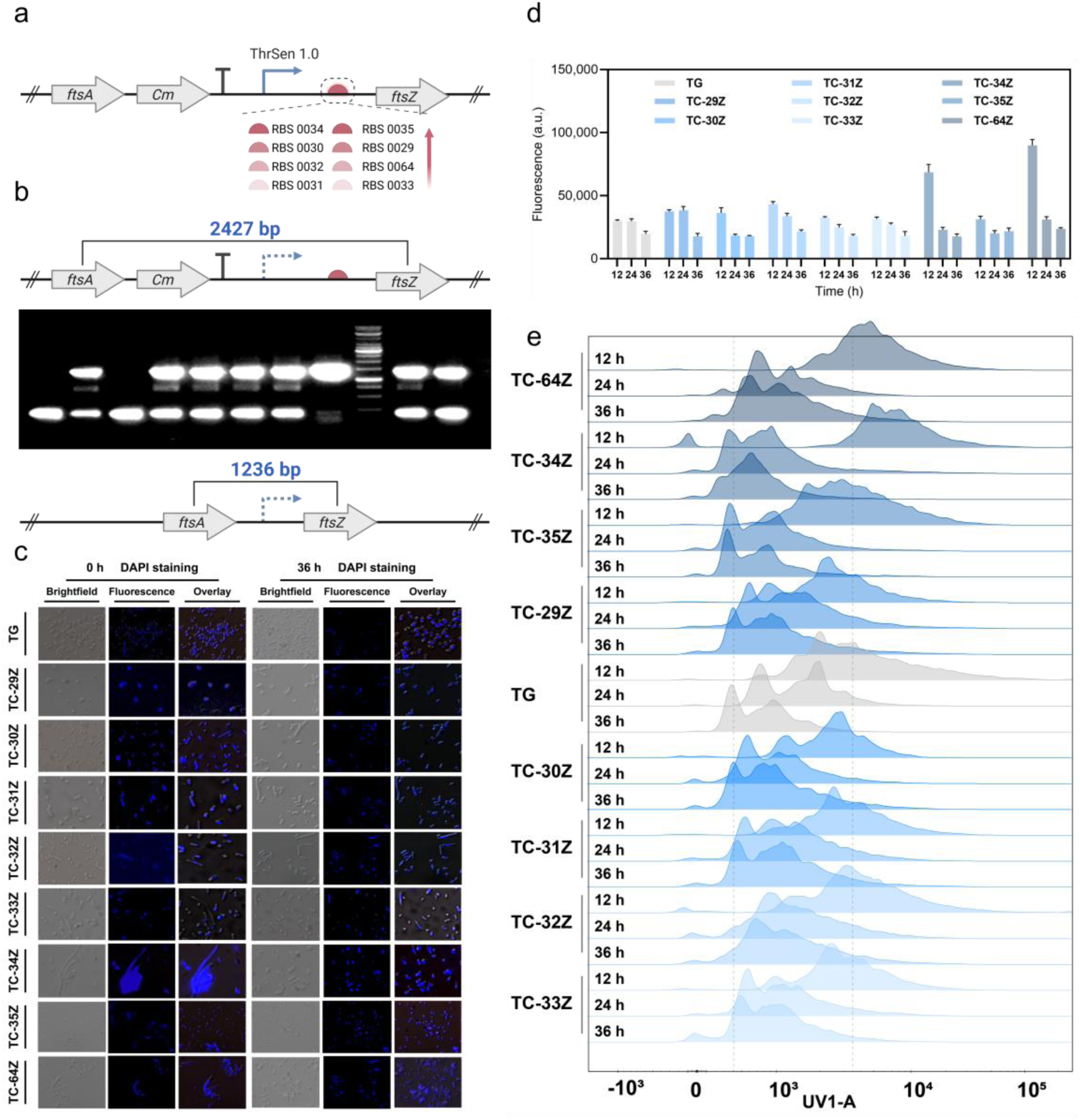
Construction and characterization of an L-threonine-responsive dynamic chromosome-ploidy control system in the production strain TG. a, Schematic of the *ftsZ* regulatory modules comprising the L-threonine-responsive element ThrSen1.0 and RBSs of different strengths, together with the proposed mechanism of chromosome-ploidy transition. b, PCR genotyping of the wild-type and engineered *ftsZ* loci in the candidate strains. c, Representative DAPI fluorescence micrographs of TG and the candidate strains at the indicated fermentation time points. d, DAPI fluorescence intensities of TG and the candidate strains quantified using a microplate reader. e, Flow-cytometric distributions of single-cell DAPI fluorescence in TG and the candidate strains, showing changes in relative cellular DNA content over the course of fermentation.

During rapid growth, when TG cells contained multiple replication forks, homologous recombination integrated the regulatory modules immediately upstream of the *ftsZ* start codon. According to the system design, intracellular L-threonine concentrations remain relatively low during the early stage of fermentation, so ThrSen1.0 weakly activates *ftsZ* on the engineered chromosome. Consequently, the engineered chromosome alone cannot support efficient cell division. Under chloramphenicol selection, cells containing both engineered and unmodified chromosome copies gain a selective advantage because they combine chloramphenicol resistance with sufficient *ftsZ* expression to sustain normal division, thereby facilitating the establishment and maintenance of a polyploid state. As fermentation progresses and intracellular L-threonine accumulates, ThrSen1.0 activity increases, enhancing *ftsZ* expression from the engineered chromosome. This progressively reduces the requirement for unmodified chromosome copies and promotes a transition toward a lower-ploidy state. Candidate strains carrying different RBSs were designated TC-29Z, TC-30Z, TC-31Z, TC-32Z, TC-33Z, TC-34Z, TC-35Z and TC-64Z (Fig. 2a).

To validate the endogenous-metabolite-driven ploidy-control system, we cultivated the candidate strains in shake flasks and collected samples at different fermentation stages. We characterized chromosome-ploidy dynamics using PCR, DAPI staining, and flow cytometry, providing complementary information on genetic composition, DNA content, and single-cell population distributions. PCR analysis using primers spanning the recombination site detected products corresponding to both the unmodified and engineered chromosome types in TC-29Z, TC-31Z, TC-32Z, TC-33Z, TC-34Z and TC-64Z (Fig. 2b). These results indicated that chromosome copies with distinct genetic configurations coexisted in these strains, providing preliminary genetic evidence for polyploid formation. Because PCR is highly sensitive to low-abundance templates and therefore cannot accurately quantify the relative abundance of the different chromosome types within a population, DAPI staining and flow cytometry were subsequently used to characterize their dynamic changes.

DAPI fluorescence imaging revealed that the candidate polyploid strains had enlarged nucleoid regions and longer cells during the early stage of fermentation, consistent with increased chromosome content. As fermentation progressed, their nucleic-acid fluorescence patterns and cellular morphologies gradually approached those of parental TG (Fig. 2c). Quantification using a microplate reader further showed that, at 12 h, the DAPI fluorescence intensities of TC-34Z and TC-64Z reached 70,224 and 91,609 arbitrary units (a.u.), respectively, markedly exceeding the 31,157 a.u. measured for TG. These results indicated that TC-34Z and TC-64Z contained substantially more DNA during the early stage of fermentation (Fig. 2d).

We next used flow cytometry to analyze DNA-content distributions at the single-cell level across different fermentation stages. At 12 h, the main fluorescence peaks of TC-34Z and TC-64Z shifted markedly toward higher intensities relative to TG, indicating higher DNA content in both cell populations. As fermentation progressed from 12 to 36 h, the fluorescence peaks of both strains gradually shifted toward lower intensities. They ultimately approached that of TG, demonstrating a progressive decline in cellular chromosome content. By contrast, although TC-29Z, TC-31Z, TC-32Z, and TC-33Z exhibited polyploid-associated features during the initial characterization, their flow-cytometric distributions at 12 h were already similar to that of TG, suggesting that their ploidy transitions occurred earlier. The distinct transition profiles of the candidate strains indicate that RBS strength may modulate the duration of the polyploid state and the timing of the transition toward lower ploidy by shaping *ftsZ* expression dynamics (Fig. 2e).

Together, the PCR, DAPI-staining, and flow-cytometric results demonstrate the establishment of an endogenous-product-responsive dynamic chromosome-ploidy control system in an L-threonine-producing strain. Among the candidate strains, TC-34Z and TC-64Z exhibited the clearest temporal transitions in DNA content. We therefore selected them for subsequent analyses of how dynamic ploidy regulation influences cellular physiology and L-threonine biosynthesis.

### Phenotypic characterization of the dynamically polyploid strain TC-64Z

Dynamic changes in chromosome ploidy alter not only the amount of intracellular genetic material but may also reshape cellular morphology and physiology^23, 24^. To further characterize phenotypes associated with dynamic polyploidy, we systematically evaluated the engineered strains for cell morphology, environmental stress tolerance, membrane permeability, cellular activity, and heterologous protein production.

We first quantified cell morphology by microscopy during the early exponential and early stationary phases. During early exponential growth, cell width remained predominantly within the range of 0.8–1.1 μm across the dynamically polyploid strains and differed little from that of the parental TG strain. By contrast, cell length increased to varying degrees, with pronounced elongation observed in TC-32Z, TC-34Z and TC-64Z (Fig. 3a–d). Accordingly, the estimated cell volume and surface area increased in the dynamically polyploid strains. TC-64Z exhibited a mean cell volume of 16.14 μm³ and a mean surface area of 30.93 μm², corresponding to 4.73-fold and 2.81-fold the respective values of TG. This cellular enlargement was accompanied by a corresponding reduction in the surface-area-to-volume ratio (Fig. 3e–i).

**Fig. 3.**
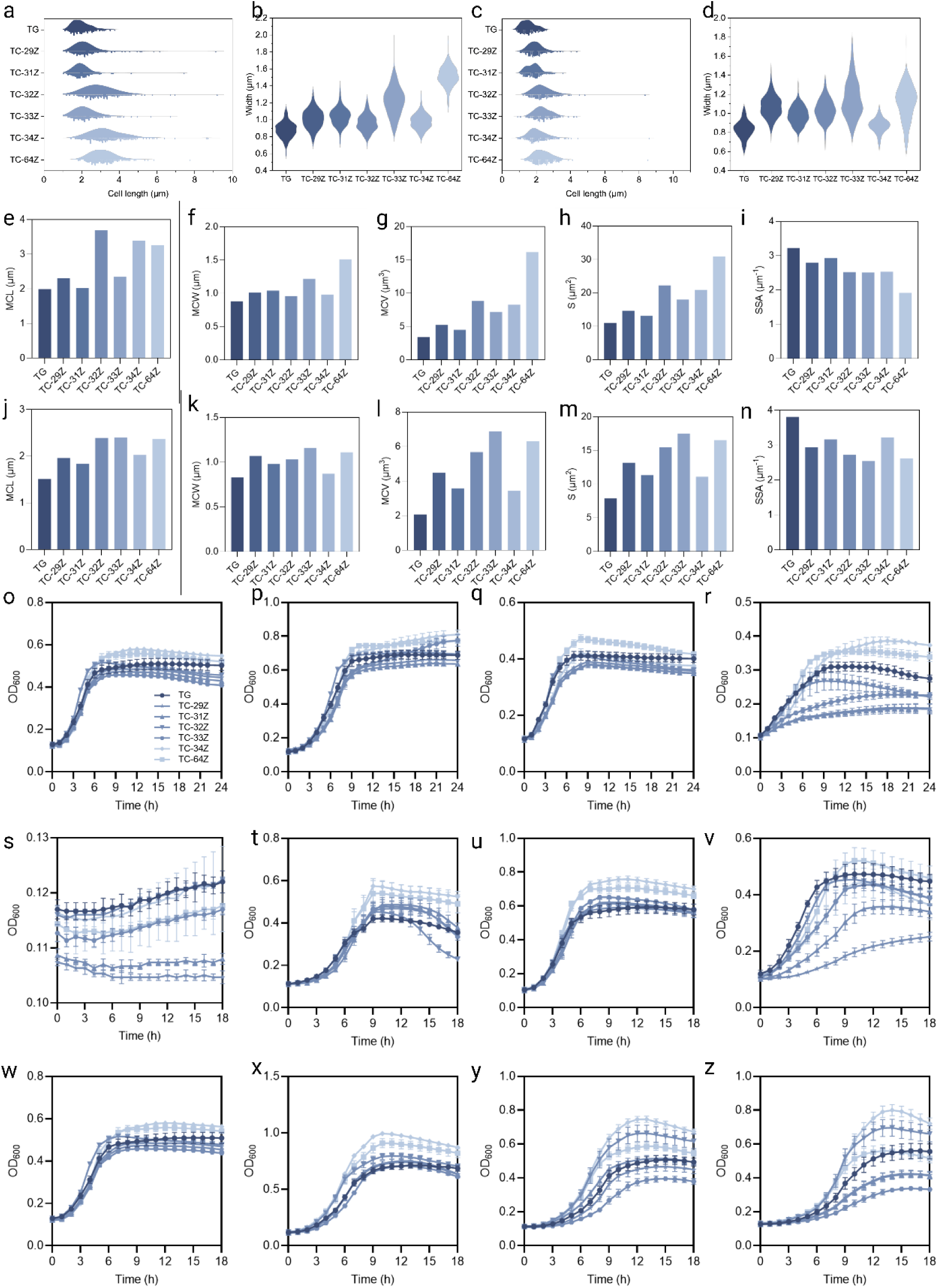
Morphological and stress-response phenotypes of dynamically polyploid strains derived from the L-threonine-producing strain TG. a,b, Violin plots showing the distributions of cell length (a) and cell width (b) during the early exponential phase. c,d, Violin plots showing the distributions of cell length (c) and cell width (d) during the early stationary phase. e–i, Mean cell length (MCL; e), mean cell width (MCW; f), mean cell volume (MCV; g), cell surface area (S; h) and specific surface area, defined as the surface-area-to-volume ratio (SSA; i), during the early exponential phase. j–n, MCL (j), MCW (k), MCV (l), S (m) and SSA (n) during the early stationary phase. o–r, Growth profiles of TG and the dynamically polyploid strains at 30 °C (o), 37 °C (p), 42 °C (q) and 45 °C (r). s–v, Growth profiles at pH 4 (s), pH 5 (t), pH 8 (u) and pH 9 (v). w–z, Growth profiles in the presence of 0 g/L (w), 1 g/L (x), 2 g/L (y) and 4 g/L (z) acetate. We monitored growth by measuring optical density at 600 nm (OD_600_).

Upon entering the early stationary phase, the cell length, volume, and surface area of most dynamically polyploid strains decreased and approached those of TG. In particular, the cell lengths of TC-29Z, TC-31Z and TC-34Z returned predominantly to the range of 1–2.5 μm (Fig. 3j–n). These phase-dependent morphological changes broadly paralleled the progressive decline in chromosome content during fermentation, indicating that dynamic changes in chromosome ploidy are accompanied by coordinated remodeling of cell morphology.

Industrial fermentation exposes production strains to multiple environmental stresses, including temperature fluctuations, pH perturbations, and the accumulation of inhibitory metabolites^25, 26^. We therefore compared the growth of the dynamically polyploid strains over 30–45 °C. At 45 °C, the growth of TC-29Z and TC-31Z was strongly inhibited, whereas TC-34Z and TC-64Z maintained higher biomass during the middle and late stages of cultivation (Fig. 3o–r). Under acidic conditions (pH 4), all strains showed strongly inhibited growth. At pH 5, 8 and 9, several dynamically polyploid strains exhibited an extended lag phase but ultimately reached relatively high biomass during the stationary phase. TC-64Z showed particularly robust growth at pH 9 (Fig. 3s–v).

Increasing acetate concentrations progressively inhibited TG growth, whereas TC-32Z, TC-34Z, and TC-64Z retained comparatively better growth performance (Fig. 3w–z). Thus, the effects of dynamic polyploidization on stress responses were strain-dependent, with TC-34Z and TC-64Z displaying improved relative growth under elevated temperature, alkaline pH and acetate stress.

Based on its pronounced chromosome-content transition, distinctive morphological phenotype, and comparatively robust growth under environmental stress, TC-64Z was selected for further physiological characterization. The parental TG strain and the static polyploid strain TH-103Z were included as controls. We first used fluorescein diacetate (FDA) staining to assess intracellular esterase activity as a proxy for cellular metabolic activity. At 12 h, TC-64Z exhibited an FDA fluorescence intensity of 10,219 a.u., approximately twice that of TG. Flow-cytometric analysis over 36 h further showed that the FDA-positive fraction of TC-64Z remained at approximately 80–90%, whereas that of TG declined to below 50% after 9 h. These results indicate that TC-64Z maintained a larger metabolically active population during prolonged cultivation (Fig. 4a, b).

**Fig. 4.**
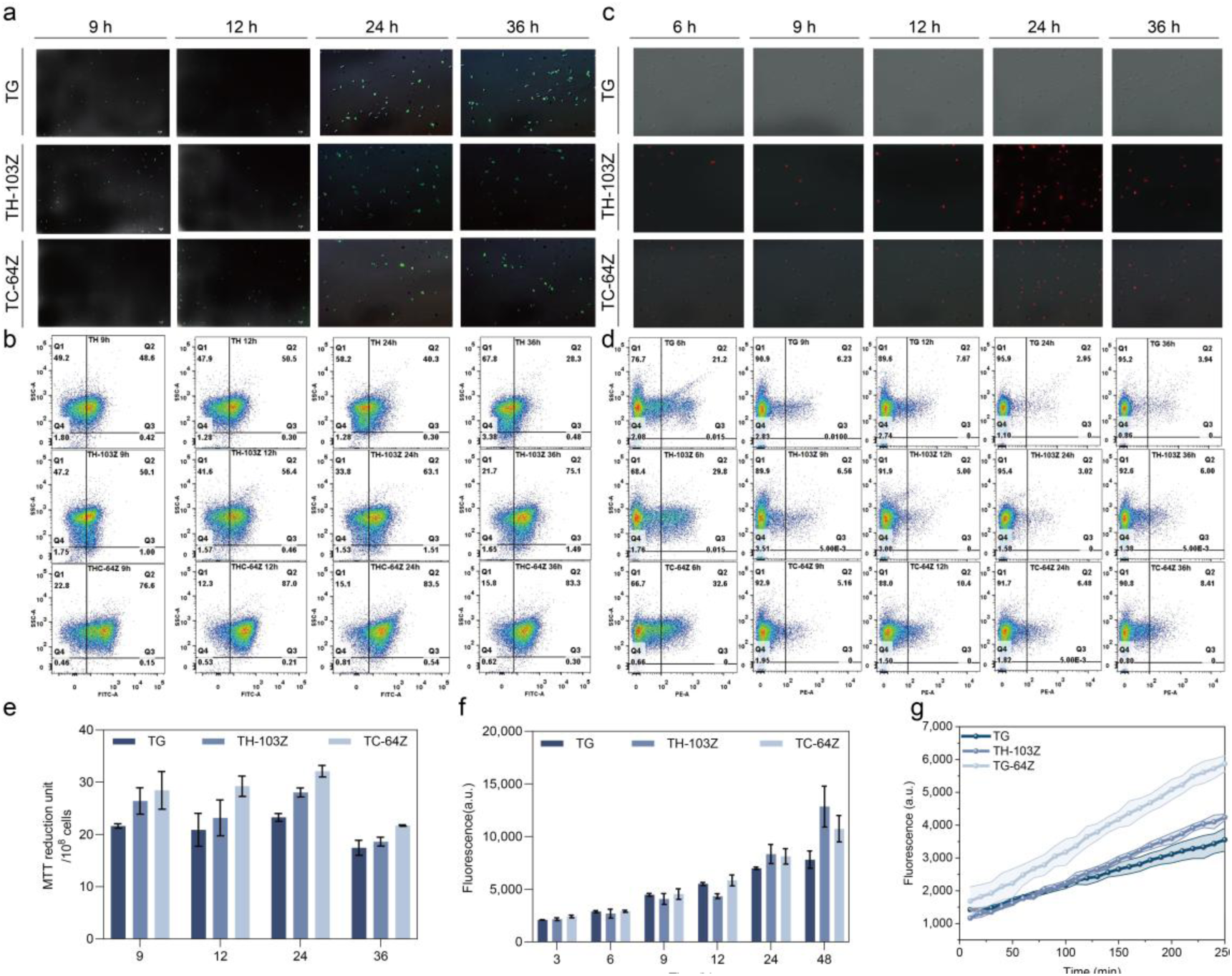
Cellular metabolic activity, membrane integrity and heterologous protein production in TC-64Z. a, Representative fluorescence micrographs of TG, TH-103Z and TC-64Z stained with fluorescein diacetate (FDA) at different cultivation times. Green fluorescence reflects intracellular conversion of FDA to fluorescein and indicates cellular esterase activity. b, Representative flow-cytometry profiles of FDA-stained TG, TH-103Z and TC-64Z cells at different cultivation times. c, Representative fluorescence micrographs of TG, TH-103Z and TC-64Z stained with propidium iodide (PI) at different cultivation times. Red fluorescence indicates PI-positive cells with increased membrane permeability or compromised membrane integrity. d, Representative flow-cytometry profiles of PI-stained TG, TH-103Z and TC-64Z cells at different cultivation times. e, Cellular metabolic activity of TG, TH-103Z and TC-64Z at different cultivation times, as determined by the MTT-reduction assay. f, Time-course RFP fluorescence of TG-RFP and TC-64Z-RFP as a proxy for heterologous protein accumulation. g, Schematic of the fluorescent nanoparticle-based assay used to evaluate M58 activity. Hydrolysis of fluorescein dilaurate (FDL) by M58 releases fluorescent fluorescein. h, Time-course fluorescence signals generated by M58 expressed in TG, TH-103Z and TC-64Z, as determined using the FDL-based activity assay.

The proportion of propidium iodide (PI)-positive cells was modestly higher in TC-64Z than in TG throughout cultivation, suggesting that dynamic polyploidization was accompanied by changes in membrane permeability or integrity (Fig. 4c,d). Consistent with the FDA results, TC-64Z also exhibited higher MTT-reduction activity than TG at all tested time points (Fig. 4e). Collectively, these findings indicate that TC-64Z maintains elevated cellular metabolic activity while displaying measurable changes in membrane physiology, providing a physiological basis for its fermentation phenotype.

Finally, we investigated whether dynamic polyploidization affected heterologous protein production. A strong constitutive promoter drove RFP expression. During the first 12 h of fermentation, fluorescence accumulation was similar between TC-64Z-RFP and TG-RFP. However, the difference between the two strains increased progressively with cultivation time. At 48 h, TG-RFP exhibited a fluorescence intensity of 7,815.87 a.u., whereas TC-64Z-RFP reached 10,748.6 a.u., representing an increase of approximately 38% over TG (Fig. 4f).

To further validate this phenotype using a functional enzyme, we expressed the plastic-degrading enzyme M58 in different chassis strains. After 4 h of catalysis, TC-64Z/M58 generated a fluorescence signal of 6,232.0 a.u., compared with 3,556.3 a.u. for the parental control, corresponding to an increase of 75.2% (Fig. 4g). Thus, in addition to its altered morphology and physiological state, TC-64Z exhibited an enhanced capacity for heterologous protein production. These findings indicate that dynamic chromosome-ploidy regulation can reshape an engineered strain’s protein-production phenotype and highlight TC-64Z’s potential as a chassis for microbial biomanufacturing.

### Dynamic polyploidy enhances L-threonine biosynthesis and fermentation performance

The results above showed that changes in cell morphology, cellular activity, and membrane properties accompanied dynamic changes in chromosome ploidy. To determine whether these phenotypic changes translated into enhanced product synthesis, we first compared the growth kinetics and L-threonine production of the parental TG strain and the dynamically polyploid strains in shake-flask fermentation. During early exponential growth, most dynamically polyploid strains grew more slowly than the TG strain. As fermentation progressed, however, the growth of TC-32Z, TC-34Z and TC-64Z gradually recovered, ultimately resulting in relatively high biomass during the late fermentation stage (Fig. 5a). This two-stage growth pattern broadly coincided with the temporal decline in chromosome content from an initially elevated state, suggesting that dynamic changes in chromosome ploidy may be associated with phase-dependent adjustments in cellular growth.

**Fig. 5.**
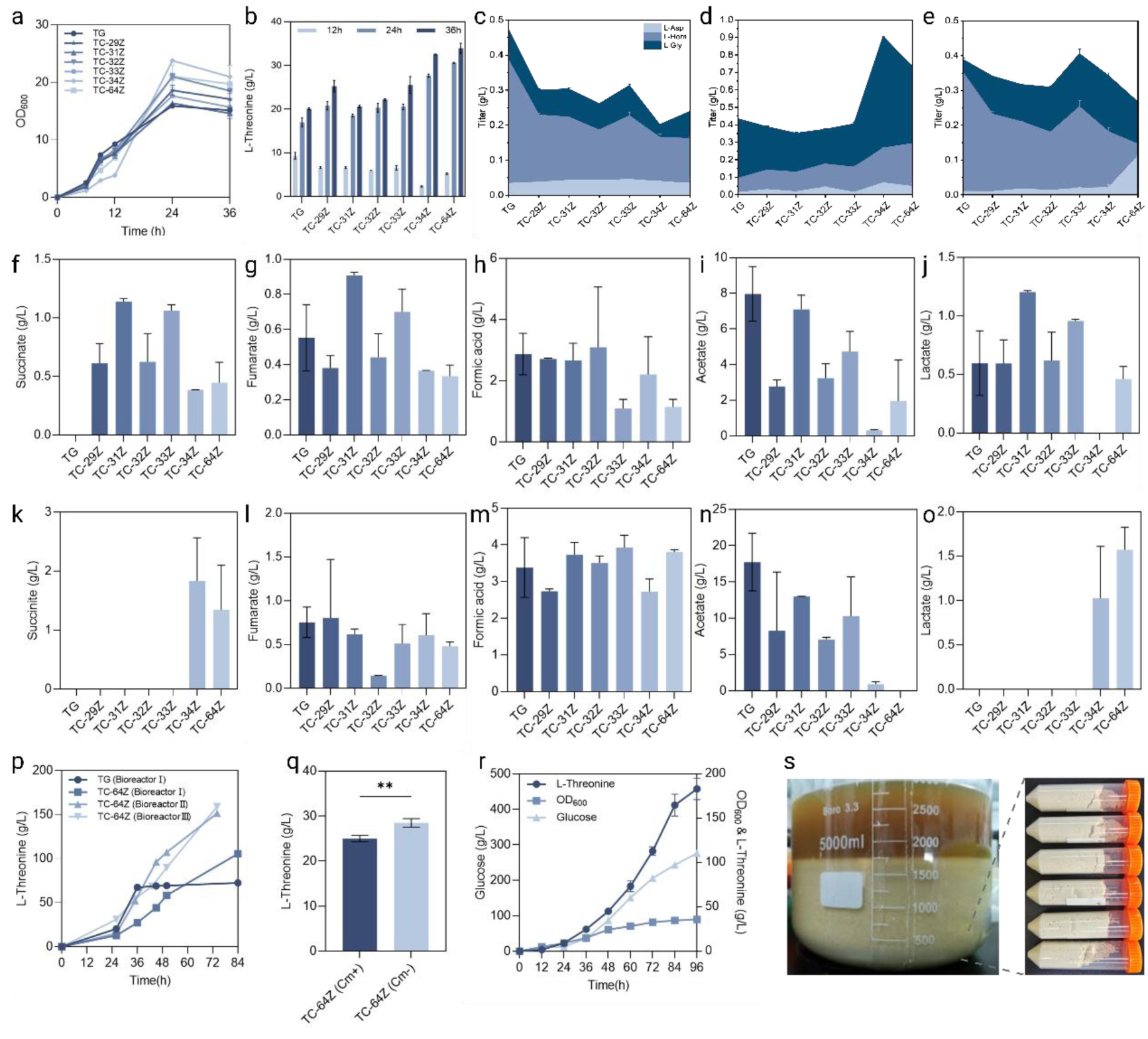
Dynamic polyploidy enhances L-threonine production in strains derived from the L-threonine-producing strain TG. a, Growth profiles of TG and the dynamically polyploid strains during shake-flask fermentation, as monitored by OD_600_. b, L-Threonine titers of TG and the dynamically polyploid strains after 12, 24 and 36 h of shake-flask fermentation. c–e, Concentrations of selected amino acid intermediates and related metabolites, including L-aspartate (L-Asp), L-homoserine (L-Hom) and glycine (Gly), after 12 h (c), 24 h (d) and 36 h (e) of shake-flask fermentation. f–j, Concentrations of succinate (f), fumarate (g), formate (h), acetate (i) and lactate (j) after 24 h of shake-flask fermentation. k–o, Concentrations of succinate (k), fumarate (l), formate (m), acetate (n) and lactate (o) after 36 h of shake-flask fermentation. p, L-Threonine production profiles of TG and TC-64Z under different 5-L fed-batch fermentation conditions. Bioreactor condition I represents the baseline fermentation process, condition II represents cultivation with an increased dissolved-oxygen level, and condition III represents the process with further optimization of agitation and aeration during the middle and late stages of fermentation. q, Final L-threonine titers of TC-64Z during shake-flask fermentation in the presence (Cm⁺) or absence (Cm⁻) of chloramphenicol. r, Time courses of residual glucose concentration, biomass accumulation (OD_600_) and L-threonine production during antibiotic-free fed-batch fermentation of TC-64Z in a 5-L bioreactor. s, Photographs of the fermentation broth after static incubation and the resulting crystalline precipitate. The white precipitate collected at the bottom was identified as L-threonine crystals.

**Fig. 6.**
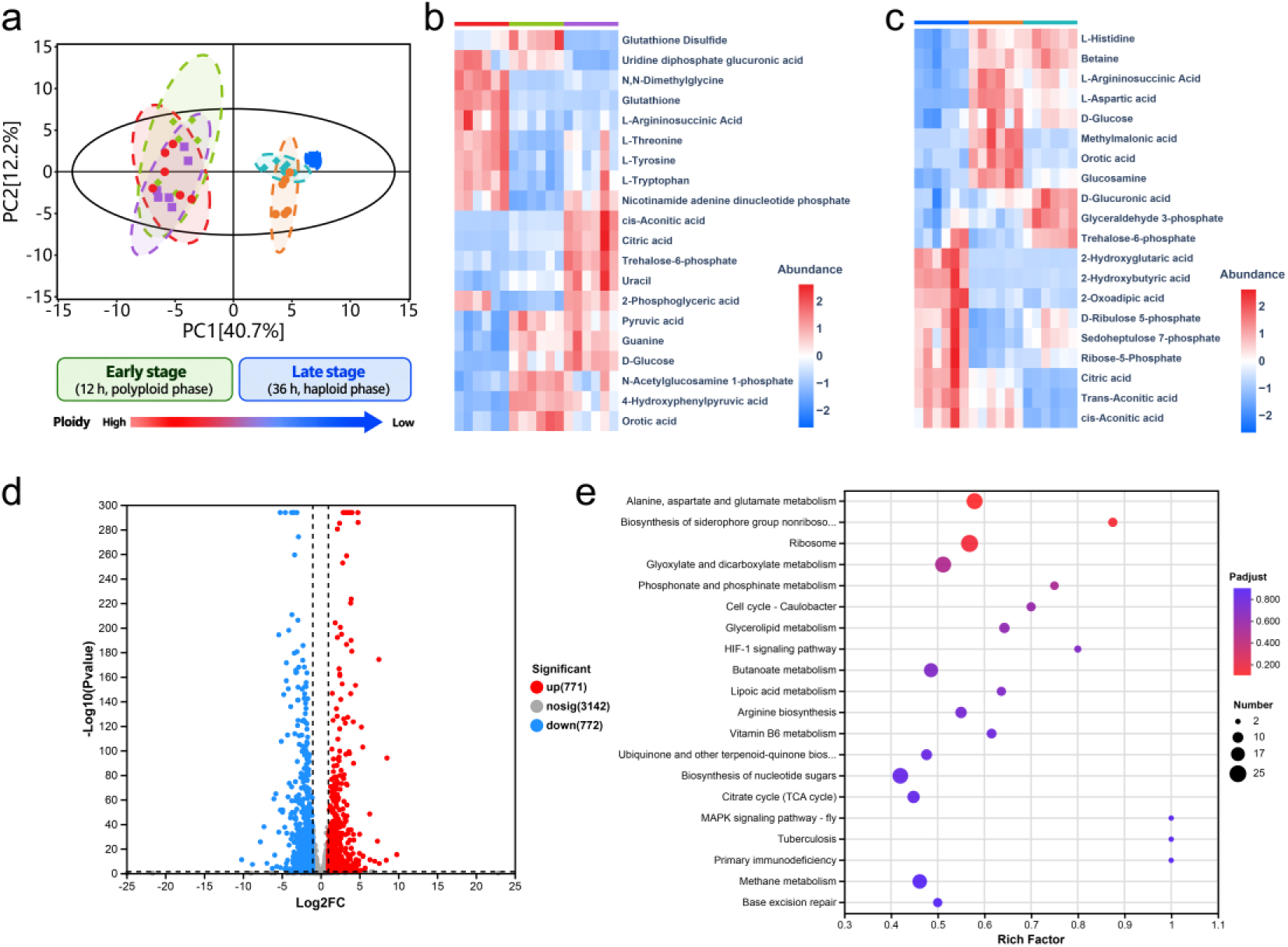
Integrated metabolomic and transcriptomic analyses reveal stage-dependent metabolic reprogramming in the dynamically polyploid strain TC-64Z. a, Principal component analysis of the metabolomic profiles of the parental strain TG, static polyploid strain TH-103Z and dynamically polyploid strain TC-64Z at 12 and 36 h of fermentation. The schematic below indicates the transition from the early high-ploidy phase to the late low-ploidy phase. b, c, Heatmaps showing the relative abundance of selected differential metabolites in TG, TH-103Z and TC-64Z at 12 h (b) and 36 h (c). Metabolite abundance was normalized by row-wise *z*-score transformation; red and blue indicate relatively high and low abundance, respectively. d, Volcano plot showing differentially expressed genes in TC-64Z relative to TG. Red and blue denote significantly upregulated and downregulated genes, respectively, whereas grey denotes genes without significant differential expression. Dashed lines indicate the thresholds used to define differential expression. e, KEGG pathway enrichment analysis of the differentially expressed genes. Dot size represents the number of differentially expressed genes assigned to each pathway, dot color indicates the adjusted *P* value, and the enrichment factor denotes the proportion of differentially expressed genes among all genes annotated to the corresponding pathway.

L-Threonine accumulation exhibited a similar temporal pattern. During the first 12 h of fermentation, the dynamically polyploid strains produced slightly less L-threonine than TG. With prolonged cultivation, however, product accumulation in these strains progressively accelerated. At 36 h, the L-threonine titer of TG was 20.04 g/L, whereas the dynamically polyploid strains exhibited increases of 10.53–69.16%. TC-64Z achieved the highest titer among all engineered strains, reaching 33.9 g/L (Fig. 5b). These results demonstrate that dynamic ploidy regulation markedly enhances the final capacity for L-threonine production.

To examine the relationship between dynamic ploidy transitions and metabolite distribution, we quantified key metabolic intermediates and by-products at different fermentation stages. At 12 h, TG accumulated 0.352 g/L L-homoserine, whereas its concentration remained below 0.2 g/L in most dynamically polyploid strains. This reduction suggests that the downstream conversion of L-homoserine may have been enhanced in the dynamically polyploid strains (Fig. 5c–e). As fermentation progressed, marked differences also emerged in metabolites associated with central carbon metabolism. At 36 h, TC-34Z and TC-64Z accumulated higher concentrations of α-ketoglutarate, succinate and lactate. In some of the high-producing strains, α-ketoglutarate and succinate reached 1.304 and 1.344 g/L, respectively, whereas lactate reached 1.571 g/L. By contrast, acetate and fumarate accumulated to lower levels than in TG, while formate showed comparatively minor changes (Fig. 5f–o). Thus, dynamic ploidy regulation affected not only the accumulation of the final L-threonine product but also the distribution of central-carbon intermediates and fermentation by-products. These observations provided a basis for subsequent transcriptomic and metabolomic analyses of the underlying metabolic regulatory mechanisms.

Following validation in shake flasks, TC-64Z was selected for fed-batch fermentation in a 5-L bioreactor to assess its performance upon scale-up. Under the baseline fermentation conditions, TG and TC-64Z achieved final L-threonine titers of 72.49 and 105.44 g/L, respectively, after 84 h. Increasing the dissolved-oxygen level during cultivation further raised TC-64Z’s L-threonine titer to 151.4 g/L. Subsequent optimization of the agitation rate and aeration during the middle and late stages of fermentation increased the titer to 158.9 g/L, with a glucose-to-L-threonine yield of 0.64 g/g and a volumetric productivity of 2.15 g/L/h (Fig. 5p). These results demonstrate efficient L-threonine production using the dynamically polyploid chassis. Importantly, TC-64Z retained its enhanced production phenotype during the transition from shake-flask cultivation to bioreactor fermentation, supporting the applicability of dynamic ploidy regulation under high-cell-density fermentation conditions.

Because continuous antibiotic supplementation increases the operational complexity of industrial fermentation, we next evaluated the dynamic polyploidy system’s production performance without antibiotics. In 12-well plate cultures, removal of the antibiotic increased the L-threonine titer of TC-64Z from 6.07 to 8.72 g/L. Similarly, in shake-flask fermentation, the final titer increased from 24.99 to 28.49 g/L (Fig. 5q). Using the optimized fermentation process described above, we subsequently performed a 96-h fed-batch fermentation in a 5-L bioreactor without antibiotic supplementation. Under these conditions, TC-64Z achieved an L-threonine titer of 183.1 g/L, a substrate-to-product conversion yield of 66.50% and a volumetric productivity of 1.907 g/L/h (Fig. 5r). Following completion of the fermentation and subsequent static incubation, pronounced crystallization of L-threonine was observed (Fig. 5s). Collectively, these results show that the dynamic polyploidy system sustains high-level L-threonine production under antibiotic-free conditions and has potential for further development as a scalable fermentation platform.

**Table 1.**
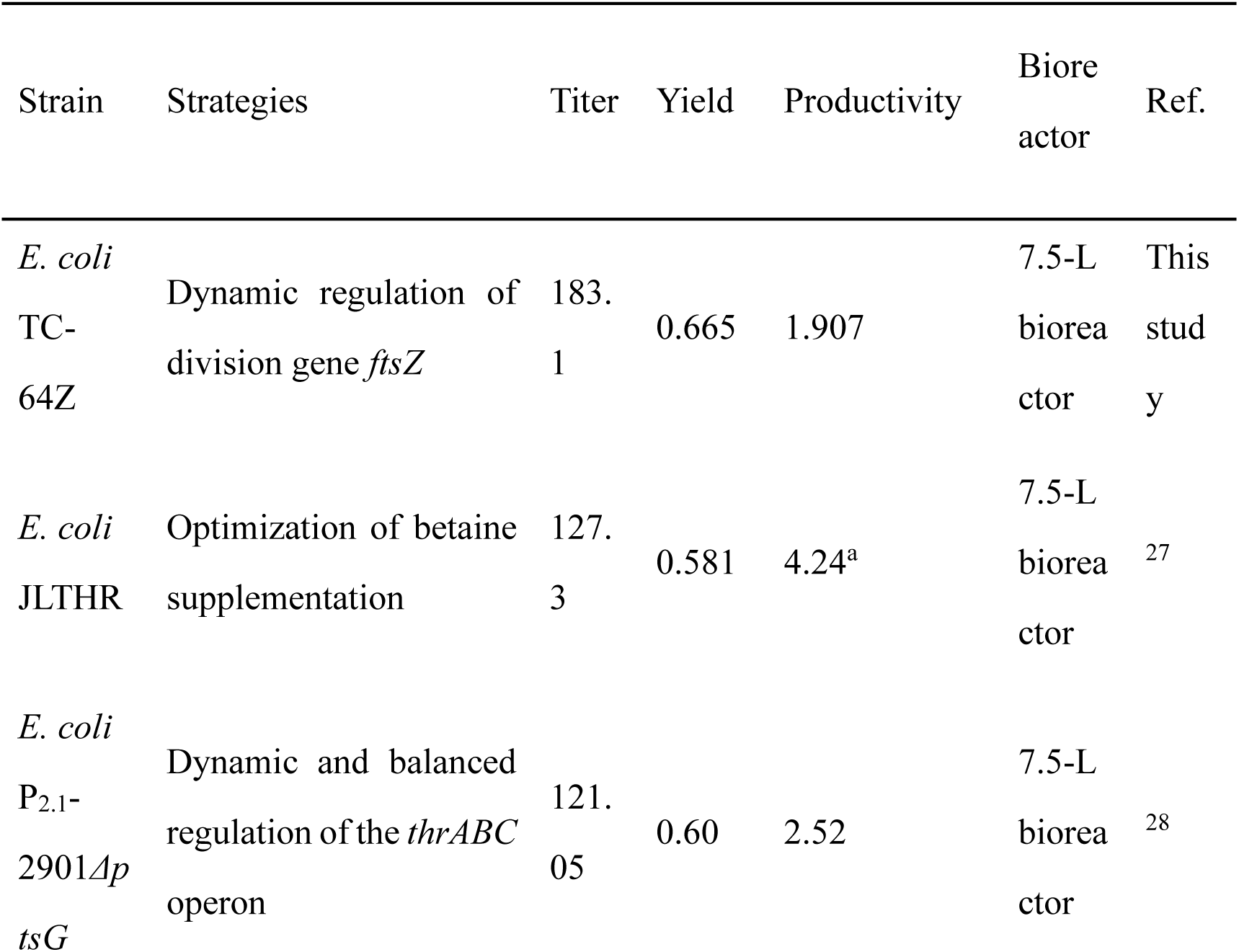
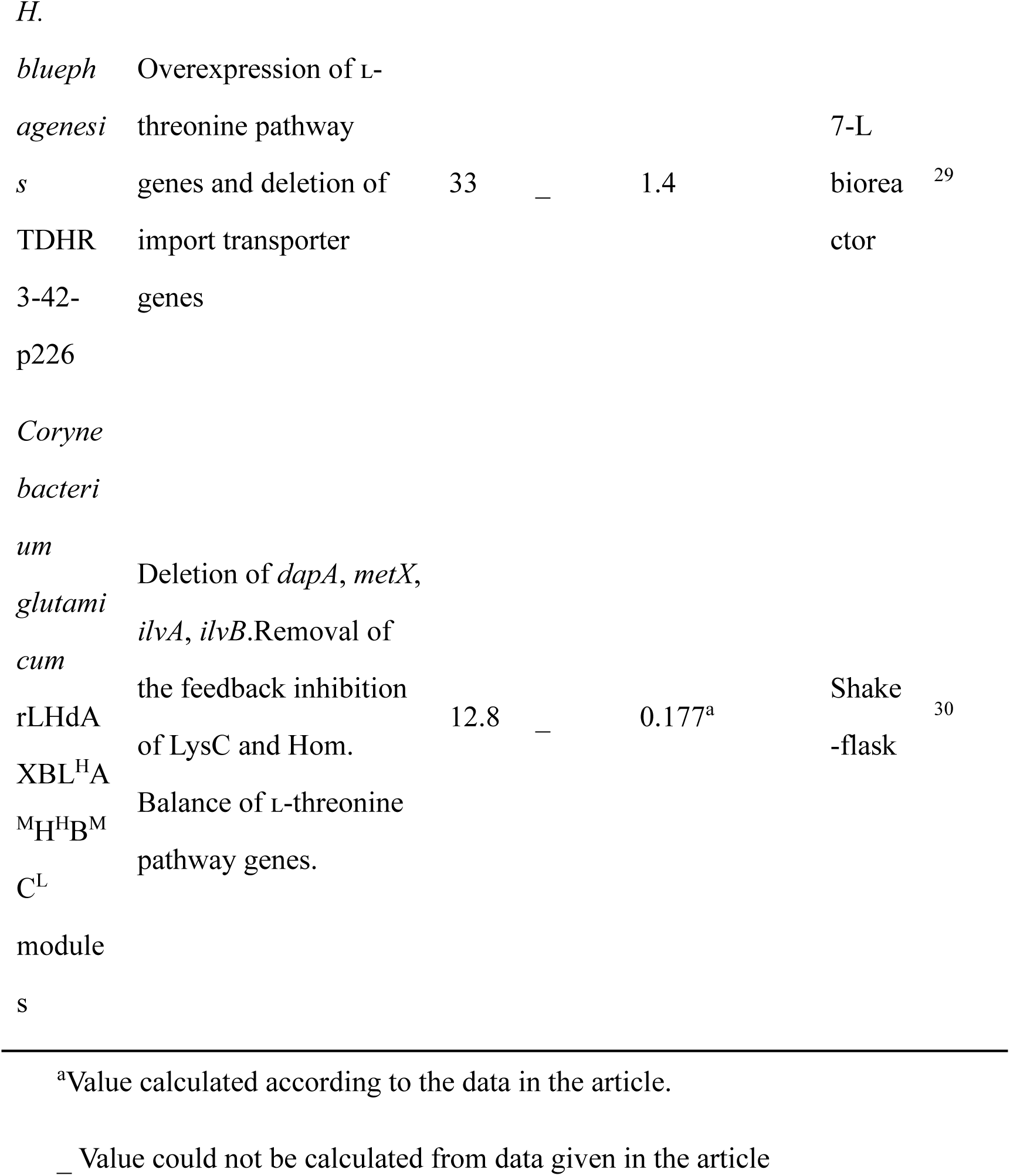
Comparison of ʟ-threonine production in various microorganisms.

### Dynamic chromosome-ploidy regulation promotes L-threonine biosynthesis by reprogramming metabolic networks

To define the metabolic basis through which dynamic chromosome-ploidy regulation enhances L-threonine biosynthesis, we compared the metabolomic profiles of the parental strain TG, the static polyploid strain TH-103Z and the dynamically polyploid strain TC-64Z at 12 and 36 h of fermentation. We further integrated these data with transcriptomic profiling of TC-64Z to examine the relationship between dynamic ploidy transitions and host metabolic-network remodeling at both the metabolite and gene-expression levels. Principal component analysis showed that PC1 and PC2 accounted for 40.7% and 12.2% of the total variance, respectively, with the six sample groups displaying a clear time-dependent distribution in metabolic space. At 12 h, TG, TH-103Z, and TC-64Z were predominantly positioned along the negative PC1 axis. In contrast, all three strains shifted markedly toward the positive PC1 axis at 36 h, indicating that fermentation progression was a major determinant of the global metabolic state. Notably, the metabolic profile of TC-64Z remained distinct from those of TG and TH-103Z at both time points, and TC-64Z formed a discrete cluster from both the parental and static polyploid strains at 36 h. Our preceding ploidy measurements showed that TC-64Z’s chromosome content had declined to near-parental levels by this stage. Nevertheless, its metabolic state did not simply revert to that of TG; instead, it developed a distinct metabolic phenotype. Thus, dynamic ploidy transitions not only alter chromosome copy number but also involve fermentation-stage-dependent remodeling of cellular metabolism.

During the high-ploidy phase at 12 h, TC-64Z already exhibited a metabolite distribution that was clearly distinguishable from those of TG and TH-103Z. The differential metabolites were primarily associated with central carbon metabolism, amino acid metabolism, redox homeostasis and nucleotide metabolism. Metabolites linked to glycolysis and the tricarboxylic acid (TCA) cycle—including citrate, cis-aconitate, pyruvate and 2-phosphoglycerate—were markedly altered. Glutathione and oxidized glutathione, metabolites associated with nicotinamide adenine dinucleotide phosphate metabolism, and uracil also underwent varying degrees of redistribution. In addition, amino acid-related metabolites, including L-threonine, L-tyrosine, L-tryptophan and L-argininosuccinate, differed substantially among the three ploidy states. Importantly, TC-64Z did not display a uniform increase in all biosynthetic products at this stage; instead, individual metabolites changed in different directions. These findings indicate that the high-ploidy state does not merely amplify global metabolism through increased gene dosage. Rather, it induces coordinated adjustments across central carbon, redox and amino acid metabolism, establishing an early, preconfigured metabolic state that may facilitate the subsequent transition to production.

By 36 h, the metabolic profile of TC-64Z had undergone further reorganization and increasingly exhibited features associated with L-threonine production. Relative to TG and TH-103Z, TC-64Z showed pronounced changes in L-aspartate, L-argininosuccinate, glyceraldehyde 3-phosphate and trehalose 6-phosphate. In contrast, TCA-cycle intermediates, including citrate, cis-aconitate and trans-aconitate, generally exhibited lower relative abundance. L-Aspartate represents the key entry point through which oxaloacetate is channelled into the biosynthesis of aspartate-family amino acids, of which L-threonine is a major end product. The concurrent enhancement of the L-aspartate-associated metabolic state and reduction in several TCA-cycle intermediates is therefore consistent with a model in which central carbon metabolism is rebalanced away from the TCA cycle and towards aspartate-family amino acid biosynthesis. Moreover, the metabolic profile of TC-64Z at 36 h remained distinct from that of the static polyploid strain TH-103Z at the same time point. These metabolic changes therefore cannot be attributed solely to an increase in chromosome copy number but are more likely associated with the dynamic ploidy transition that occurs over the course of fermentation.

Transcriptomic analysis further supported this metabolic-network remodeling at the gene-expression level. Using the parental TG strain as the reference, we identified 1543 significantly differentially expressed genes in TC-64Z: 771 upregulated and 772 downregulated. KEGG enrichment analysis showed that these genes were broadly distributed across pathways involved in alanine, aspartate, and glutamate metabolism; glyoxylate and dicarboxylate metabolism; the TCA cycle; pyruvate metabolism; ribosome biogenesis; and cell-cycle-related processes. Thus, dynamic ploidy regulation was not confined to the L-threonine biosynthetic pathway but encompassed systematic adjustments to central metabolism and fundamental cellular physiology. Further examination of amino acid metabolism revealed increased expression of multiple genes involved in L-threonine and aspartate-family amino acid biosynthesis. In particular, *thrA* and *lysC* were upregulated by 11.362-fold and 6.862-fold, respectively, whereas TCA-cycle-associated genes exhibited an overall downward trend. These transcriptional changes mirrored the enhanced aspartate-associated metabolic state and reduced abundance of citrate and aconitate intermediates observed in the 36-h metabolome. Together, the transcriptomic and metabolomic data support a coordinated remodeling of central carbon metabolism and the aspartate–L-threonine biosynthetic network during the dynamic ploidy transition.

Integrating the time-resolved metabolomic and transcriptomic data, we propose a stage-specific model of metabolic reprogramming driven by dynamic chromosome ploidy. During the early polyploid phase of fermentation, elevated chromosome content is accompanied by extensive changes in central carbon metabolism, redox metabolism, nucleotide metabolism, and multiple amino acid pathways, establishing a metabolic state distinct from both the parental strain and the static polyploid strain. As L-threonine accumulates and triggers a progressive decline in chromosome content, the cellular metabolic network shifts toward a production-oriented state characterized by an attenuated TCA-cycle-associated signature and reinforced metabolic nodes supporting aspartate-family amino acid and L-threonine biosynthesis. The static polyploid strain TH-103Z did not exhibit the same time-dependent metabolic transition as TC-64Z, suggesting that dynamic ploidy switching, rather than the sustained maintenance of a high chromosome copy number, is a key determinant of this metabolic-state transition. Dynamic chromosome-ploidy regulation may therefore temporally coordinate gene dosage, central carbon metabolism, and pathway-specific demand, enabling the host to progress from an early high-ploidy metabolic state to a production state favourable for L-threonine accumulation. This model provides a multi-omics mechanistic framework for TC-64Z’s high-production phenotype.

## Discussion

In this study, we established a dynamic chromosome-ploidy control system driven by intracellular metabolic state. We used L-threonine production as a model to examine the system-level effects of temporal changes in chromosome ploidy on cellular physiology, metabolic networks and production performance. By coupling an L-threonine-responsive regulatory element to the essential cell-division gene *ftsZ*, the engineered strains progressively transitioned from a high-ploidy state to a low-ploidy state as L-threonine accumulated during fermentation. PCR-based analyses, DAPI staining and flow cytometry provided complementary evidence for this dynamic process. In particular, TC-34Z and TC-64Z showed elevated DNA content during early fermentation, which subsequently declined toward that of the haploid control as fermentation progressed. Importantly, this ploidy transition was not an isolated genetic phenomenon; stage-specific changes in cell morphology, physiological activity, and L-threonine production accompanied it. These findings establish chromosome ploidy as an engineerable cellular state variable that can change over the course of fermentation and be functionally coupled to production.

The production advantage of the dynamically polyploid strains cannot be explained solely by the gene-dosage effect arising from increased chromosome copy number. Time-resolved metabolomic profiling revealed distinct metabolic states among the haploid strain TG, the static polyploid strain TH-103Z and the dynamically polyploid strain TC-64Z at both 12 and 36 h. Most notably, TC-64Z showed a distinct metabolic phenotype that differed from TG and TH-103Z at 36 h. At this stage, TC-64Z’s chromosome content had already declined to near-haploid levels, yet its metabolic state did not concomitantly revert to the original haploid state represented by TG. This observation suggests that the biological effects of dynamic ploidy depend not only on chromosome copy number at a given time point but also on the preceding ploidy trajectory and the system-wide adaptations elicited during the transition. In other words, maintaining a constitutively high ploidy and undergoing a high-to-low ploidy transition do not generate equivalent cellular states. The biphasic changes in growth and product accumulation observed during shake-flask fermentation were temporally consistent with this interpretation: the dynamically polyploid strains initially grew more slowly, whereas both growth and L-threonine accumulation increased as chromosome content declined. Although these findings support stage-specific coordination between dynamic ploidy, cellular growth, and product biosynthesis, the energetic and material costs of chromosome replication—including ATP, deoxyribonucleoside triphosphates, and other cellular resources—were not directly quantified. The underlying mechanisms of resource allocation therefore remain unresolved.

The transcriptomic and metabolomic data further support the notion that dynamic ploidy transitions are accompanied by reprogramming of central carbon metabolism and amino acid biosynthesis. During the high-ploidy phase at 12 h, TC-64Z already exhibited metabolite distributions associated with glycolysis, the tricarboxylic acid (TCA) cycle, redox metabolism and multiple amino acid pathways that differed from those of TG and TH-103Z. These changes did not reflect a uniform increase across all metabolites; instead, individual metabolic nodes responded in different directions. Thus, an increase in chromosome copy number does not simply amplify the entire metabolic network in direct proportion to gene dosage; instead, it promotes coordinated adjustments among distinct metabolic processes. By 36 h, the metabolic state of TC-64Z had reorganized further. Metabolic nodes associated with central carbon metabolism and aspartate-family amino acid biosynthesis, including L-aspartate and glyceraldehyde 3-phosphate, were markedly altered.

In contrast, TCA-cycle-associated metabolites such as citrate, cis-aconitate and trans-aconitate generally displayed lower relative abundances. Consistent with these metabolic changes, differentially expressed genes in TC-64Z were enriched in pathways related to alanine, aspartate, and glutamate metabolism; the TCA cycle; glyoxylate and dicarboxylate metabolism; pyruvate metabolism; ribosome function; and the cell cycle. The key L-threonine-and aspartate-family-biosynthetic genes *thrA* and *lysC* were upregulated by 11.362-fold and 6.862-fold, respectively, whereas genes associated with the TCA cycle exhibited an overall downward trend. The concordance between the transcriptional and metabolic profiles supports a rebalancing of central carbon metabolism towards the aspartate–L-threonine biosynthetic network during the dynamic ploidy transition.

Pronounced changes in cell morphology and physiological state also accompanied dynamic ploidy transitions. During the high-ploidy phase, the cell volume of TC-64Z reached up to 4.73 times that of the haploid TG strain, together with corresponding changes in cell-surface area and surface-area-to-volume ratio. As chromosome content declined, cellular morphology progressively approached that of the haploid state, revealing stage-specific morphological plasticity that closely tracked the ploidy transition. Fluorescein diacetate, propidium iodide and MTT assays further indicated that TC-64Z maintained a relatively high proportion of metabolically active cells during fermentation while undergoing detectable changes in membrane properties. Moreover, TC-64Z outperformed the haploid control in the expression of both red fluorescent protein and the heterologous protein M58, indicating that the effects of dynamic ploidy on cellular protein-synthesis capacity were not restricted to enzymes involved in L-threonine metabolism. Collectively, these findings suggest that the production advantage conferred by dynamic ploidy arises from coordinated changes in gene dosage, physiological state and metabolic-network organization rather than from the enhancement of a single metabolic node. Nevertheless, propidium iodide staining provides evidence only for changes in membrane integrity or permeability and does not demonstrate increased L-threonine transport across the membrane. Similarly, establishing a direct causal relationship between ploidy transitions and protein-synthesis capacity will require more targeted molecular evidence.

From an application perspective, the dynamic ploidy system exhibited robust production performance and scalability in an L-threonine-producing strain. TC-64Z produced 33.9 g/L L-threonine in shake flasks and retained strong production capacity during 5-L fed-batch fermentation. More importantly, the system uses intracellular L-threonine as an endogenous regulatory signal, thereby coupling chromosome-ploidy transitions directly to product accumulation without requiring an exogenous inducer. After removing antibiotics from the fermentation process, the engineered strain maintained high L-threonine production. It ultimately achieved a titer of 183.1 g/L together with high substrate-conversion efficiency. This endogenous metabolite-driven mode of cellular-state control reduces dependence on external process inputs and illustrates the potential of coupling chromosome-scale dynamic regulation to industrial fermentation. Although the design principle was initially established and validated in an *Escherichia coli* L-threonine production system, its underlying mechanism does not depend on the direct regulation of a single metabolic enzyme or a localized pathway. Instead, it alters chromosome state to reshape global cellular physiology and metabolism, providing a conceptual and engineering foundation for its extension to other microbial production platforms. Validation across additional microbial hosts and target products will be required to define the scope and generality of this strategy. In parallel, higher-resolution time-series multi-omics, ^13^C-based metabolic flux analysis, and direct measurements of ATP, NAD(P)H, and nucleotide pools will help clarify the causal relationships among gene-dosage variation, replication-associated resource demand, and metabolic-network remodeling. Such analyses should also inform the optimization of metabolite-sensing thresholds and the timing of ploidy transitions.

Together, our findings show that dynamic chromosome-ploidy regulation can couple target-product accumulation to changes in cellular genome state while driving stage-specific remodeling of cell morphology, physiological activity and metabolic organization. In contrast to the haploid and static polyploid strains, the dynamically polyploid strain developed a distinct and strongly time-dependent metabolic state during fermentation. Notably, convergence in chromosome content did not lead to a corresponding convergence in metabolic state, suggesting that a dynamic ploidy transition induces system-wide adaptations that continue to influence the subsequent production phase. Multi-omics analyses further support a rebalancing between central carbon metabolism and aspartate-family amino acid biosynthesis that coincides with the pronounced enhancement of L-threonine production. The significance of dynamic ploidy regulation therefore lies not in sustaining a high chromosome copy number but in using the temporal trajectory of chromosome state to shape cellular physiology and metabolism at different stages of fermentation. In this way, chromosome ploidy can be transformed from a relatively static genetic property into a metabolically responsive engineering variable that connects gene dosage, cellular state, metabolic-network organization and product biosynthesis at the chromosome scale.

## Methods

### Strains, plasmids and culture conditions

We used the industrial L-threonine-producing strain TG, maintained in our laboratory, and the model strain *Escherichia coli* MG1655 as parental strains to construct externally inducible and L-threonine-responsive dynamic chromosome-ploidy control strains, respectively.

For construction in MG1655, an IPTG-inducible expression cassette comprising FRT–*CmR*–FRT–B1006–constitutive promoter–*lacO*–RBS was integrated upstream of the start codon of the chromosomal *ftsZ* gene to enable IPTG-dependent regulation of *ftsZ* expression. The B1006 transcriptional terminator, the constitutive promoters J23109 and J23116, and the ribosome-binding sites (RBSs) B0029, B0030, B0031, B0032, B0033, B0034, B0035 and B0064, which span a range of translation strengths, were obtained from the iGEM Registry of Standard Biological Parts. Combinations of different promoters and RBSs generated the MG1655-derived strains MG-929Z, MG-930Z, MG-931Z, MG-932Z, MG-933Z, MG-934Z, MG-935Z and MG-964Z.

For construction in the L-threonine-producing strain TG, a ThrSen 1.0-based L-threonine-responsive expression cassette comprising FRT–*CmR*–FRT–B1006–ThrSen1.0–RBS was integrated upstream of the start codon of the chromosomal *ftsZ* gene. ThrSen1.0 is an L-threonine biosensor that converts intracellular L-threonine levels into a transcriptional output, thereby enabling L-threonine-dependent regulation of *ftsZ* expression. We incorporated the same series of RBSs used in the MG1655 system downstream of ThrSen 1.0. Modulating *ftsZ* translation by varying RBS strength generated strains TC-29Z, TC-30Z, TC-31Z, TC-32Z, TC-33Z, TC-34Z, TC-35Z, and TC-64Z.

We isolated candidate recombinant colonies using the corresponding antibiotic selection and verified them by PCR amplification with primers spanning the genomic integration site. The strains, plasmids, primers and regulatory elements used in this study are listed in Supplementary Tables 1–3.

### DAPI staining

Cellular DNA in TG and the dynamically polyploid strains was fluorescently labelled with 4′,6-diamidino-2-phenylindole (DAPI) during shake-flask fermentation. Samples were collected periodically during fermentation, and cells were harvested by centrifugation at 12000 rpm for 2 min. The cell pellets were washed with phosphate-buffered saline (PBS), resuspended in 1 mL of 3.7% formaldehyde and fixed at 4 °C for 40 min. The fixed cells were washed three times with PBS, resuspended in 5% Triton X-100 and incubated at 4 °C for 40 min to permeabilize the cell membrane.

The treated cells were gently washed three additional times with PBS. After centrifugation and removal of the supernatant, the cells were stained with DAPI at a final concentration of 5 mg/L for 10 min at room temperature in the dark. Remove unbound DAPI by washing the cells three times with PBS. Finally, the cells were resuspended in PBS for subsequent fluorescence microscopy, fluorescence-intensity measurements or flow-cytometric analysis.

### FDA and PI staining

We cultured strain TG and the dynamically polyploid strains in shake flasks and collected cells at the indicated time points for separate propidium iodide (PI) and fluorescein diacetate (FDA) staining analyses.

For PI staining, cell samples were washed twice with phosphate-buffered saline (PBS) and resuspended in 200 μL of PI staining solution at a concentration of 50 μg/mL. Incubate samples at 4 °C for 10 min in the dark. After staining, the cells were washed twice with PBS to remove unbound PI, resuspended in 200 μL of PBS and stored at −20 °C until analysis.

For FDA staining, wash cell samples twice with PBS, then resuspend in 200 μL of 0.05% FDA staining solution. Samples were incubated at 37 °C for 15 min in the dark, washed twice with PBS, resuspended in 200 μL of PBS and stored at −20 °C until analysis. The stained samples were subsequently analyzed by fluorescence microscopy or flow cytometry to assess cellular metabolic activity and changes in membrane integrity.

### Shake-flask fermentation for L-threonine production

A single colony was picked from an LB agar plate and inoculated into 5 ml of LB broth. The culture was incubated at 37 °C for 12 h to prepare the seed culture. The seed culture was then transferred at an inoculum ratio of 1% (v/v) into a 300-mL baffled flask containing 20 mL of fermentation medium and cultivated at 37 °C and 220 rpm for 36 h.

The glucose concentration in the medium was monitored periodically throughout fermentation. When the glucose concentration decreased below 10 g/L, glucose was supplemented to a final concentration of 40 g/L to maintain the carbon supply for continued fermentation.

### Fed-batch fermentation for L-threonine production

Frozen strain stocks maintained at −80 °C were streaked to obtain single colonies. A single colony was inoculated into a 300-ml shake flask containing 40 ml of LB broth and cultivated at 37 °C and 220 rpm for 12 h to prepare the primary seed culture. The primary seed culture was subsequently transferred at an inoculum ratio of 10% (v/v) into 400 ml of secondary seed medium and cultivated at 37 °C and 220 rpm until the exponential growth phase.

The secondary seed culture was inoculated at 10% (v/v) into a 7.5-L bioreactor containing 4 L of fed-batch fermentation medium, and fermentation was conducted at 37 °C. The pH was maintained at 7.0 by adding aqueous ammonia. During the first 12 h, the agitation speed and aeration rate were maintained at 200 rpm and 2 vvm, respectively. After 12 h, the agitation speed was increased to 1000 rpm, and the aeration rate to 10 vvm, and these conditions were maintained until the end of fermentation.

The glucose concentration in the culture broth was monitored periodically. When the glucose concentration fell below 10 g/L, an 80% (w/v) glucose solution was added using a feeding pump to restore it to 40 g/L.

### Measurement of biomass, glucose and L-threonine

Fermentation samples were collected periodically to determine cell biomass and glucose and L-threonine concentrations. To measure biomass, an appropriate volume of fermentation broth was treated with 1 M hydrochloric acid to dissolve the calcium carbonate in the medium. After dilution to an appropriate concentration, the absorbance at 600 nm (OD_600_) was measured using a UV–visible spectrophotometer as an indicator of cell biomass.

To measure glucose, we centrifuged fermentation samples to remove cells. The resulting supernatants were passed through a 0.22-μm hydrophilic membrane filter and analysed by high-performance liquid chromatography (HPLC).

To measure L-threonine, dilute the fermentation supernatant 20-fold. A 200-μl aliquot of the diluted sample was mixed with 100 μL of L-threonine derivatization reagent A and 100 μL of L-threonine derivatization reagent B and incubated at room temperature for 1 h. Then, add 400 μL of *n*-hexane, vortex for 30 s, and allow the mixture to stand at room temperature for 10 min. A 200-μL aliquot of the lower aqueous phase was transferred to a 1.5-mL microcentrifuge tube, diluted with 800 μL of deionized water, and mixed thoroughly. The diluted sample was passed through a 0.22-μm organic-solvent-compatible membrane filter and transferred to an HPLC vial.

L-Threonine was quantified using an HPLC system equipped with a Venusil AA amino acid analysis column (4.6 × 250 mm, 5-μm particle size) and an SPD detector. Chromatographic separation was performed using mobile phases A and B at a flow rate of 1.0 mL min⁻¹. The column temperature was maintained at 40 °C, the detection wavelength was set to 254 nm, and the injection volume was 10 μL.

### Transcriptome analysis

Single colonies of strains TG and TC-64Z were inoculated into 10 mL of LB medium and cultured at 37 °C for 12 h. The seed cultures were then transferred at an inoculum size of 2% (v/v) into 300 mL shake flasks containing 20 mL of shake-flask fermentation medium and cultivated at 37 °C. To maintain the polyploid state of strain TC-64Z, chloramphenicol (34 μg/mL) was added to its culture. Cells of TG and TC-64Z were collected at the mid-exponential phase for transcriptome analysis performed at OE Biotech Co., Ltd. (Shanghai, China). Total RNA was extracted using the mirVana miRNA isolation kit (Ambion). RNA integrity was assessed using an Agilent 2100 Bioanalyzer (Agilent Technologies, Santa Clara, CA, USA). Samples with RNA integrity numbers ≥ 7 were subsequently analyzed. Libraries were constructed using TruSeq Stranded Total RNA with RiboZero Gold following the manufacturer’s instructions. These libraries were then sequenced on an Illumina sequencing platform (HiSeqTM 2500) to generate 150 bp/125 bp paired-end reads. Three independent biological replicates were performed for strains TH and TC-64Z.

### Analysis of Energy Metabolome

Strains TG, TH-103Z, and TC-64Z were cultured in 300 mL shake flasks containing 20 mL of L-threonine fermentation medium at 37 °C with agitation at 220 rpm. Cells were harvested at two distinct ploidy stages: 12 h (polyploid stage) and 36 h (haploid stage). The harvested cultures were centrifuged to obtain cell pellets. Pellets were washed twice with phosphate-buffered saline (PBS), followed by immediate quenching in liquid nitrogen to arrest metabolic activity. Analysis of the energy metabolome was performed by Shanghai Biotree Biotech Co., Ltd. In brief, the sample was diluted with 300 µL water and vortex mixed for 30 s. Two steel beads were added, followed by vortex mixing for 30 s and homogenization by grinding at 35 Hz for 4 min. The sample was then sonicated for 5 min in an ice-water bath. This combined grinding and sonication cycle was repeated twice. After centrifugation, 250 µL of supernatant was mixed with 750 µL extraction solvent, vortexed for 30 s, and incubated at −40 °C for 1 h. Following centrifugation at 12 000 rpm and 4 °C for 15 min, 900 µL of supernatant was transferred to a 2 mL microcentrifuge tube and dried under vacuum. The residue was reconstituted in 180 µL of a methanol/acetonitrile/water mixture (1:1:2, v/v/v), filtered, and analyzed by UHPLC-MS/MS.

## Statistical analysis

Results are presented as the mean ± standard error of the mean. Differences between the means were evaluated using a one-way analysis of variance, with p < 0.05 was considered statistically significant.

## Acknowledgments

We thank Haiyan Yu, Xiaomin Zhao, Yuyu Guo, Xiangmei Ren, and Sen Wang of the Core Facilities for Life and Environmental Sciences, State Key Laboratory of Microbial Technology of Shandong University, for flow cytometry, HPLC, and fluorescence microscopy analyses.

## Data availability

All data collected and analyzed in this study are available from the corresponding author upon reasonable request.

## Author Contributions

X.J. and Y.G. conceived the concept. X.J. and H.S. designed the experiments. X.J., Y.G., X.Z. and X.X. designed and built the experimental platform. X.J., Y.G., H.S., X.Z., and X.X. performed the experiments. X.J. carried out the data analysis. X.J. and S.W. wrote the manuscript. S.W., Q.Q. and Q.L. provided guidance and supervised the project. All authors reviewed and approved the final manuscript.

