## Supplemental Table 1, 2, 3, and Figure 1 for "Dynamic Control of Prokaryotic Chromosome Ploidy Rewires Metabolic Networks to Enhance Product Biosynthesis"

**Supplemental Information**

**Table S1. Strains used in this study.**

| Strains | Description | Source |
|---|---|---|
| *E. coli* DH5α | $F^-$ supE44 *ΔlacU169* (*ϕ80 lacZ ΔM15*) *hsdR*17 *recA*1 *endA*1 *gyrA96 thi*-1 *relA*1 | Invitrogen |
| MG1655 | K12 F- *lambda- ilvG- rfb-50 rph-1* | Invitrogen |
| TH-103Z | L-threonine producing *E. coli* strain | Lab Stock |
| TG | L-threonine producing *E. coli* strain | Lab Stock |
| MG-929Z | Inserting DNA sequence FRT-*CmR*-FRT-B1006-J23109-*lacO*-B0029 before start codon of *ftsZ* in MG1655 | This study |
| MG-930Z | Inserting DNA sequence FRT-*CmR*-FRT-B1006-J23109-*lacO*-B0030 before start codon of *ftsZ* in MG1655 | This study |
| MG-931Z | Inserting DNA sequence FRT-*CmR*-FRT-B1006-J23109-*lacO*-B0031 before start codon of *ftsZ* in MG1655 | This study |
| MG-932Z | Inserting DNA sequence FRT-*CmR*-FRT-B1006-J23109-*lacO*-B0032 before start codon of *ftsZ* in MG1655 | This study |
| MG-933Z | Inserting DNA sequence FRT-*CmR*-FRT-B1006-J23109-*lacO*-B0033 before start codon of *ftsZ* in MG1655 | This study |

| | | |
|---|---|---|
| MG-934Z | Inserting DNA sequence FRT-*CmR*-FRT-B1006-J23109-*lacO*-B0034 before start codon of *ftsZ* in MG1655 | This study |
| MG-935Z | Inserting DNA sequence FRT-*CmR*-FRT-B1006-J23109-*lacO*-B0035 before start codon of *ftsZ* in MG1655 | This study |
| MG-964Z | Inserting DNA sequence FRT-*CmR*-FRT-B1006-J23109-*lacO*-B0064 before start codon of *ftsZ* in MG1655 | This study |
| MG-629Z | Inserting DNA sequence FRT-*CmR*-FRT-B1006-J23116-*lacO*-B0029 before start codon of *ftsZ* in MG1655 | This study |
| MG-630Z | Inserting DNA sequence FRT-*CmR*-FRT-B1006-J23116-*lacO*-B0030 before start codon of *ftsZ* in MG1655 | This study |
| MG-631Z | Inserting DNA sequence FRT-*CmR*-FRT-B1006-J23116-*lacO*-B0031 before start codon of *ftsZ* in MG1655 | This study |
| MG-632Z | Inserting DNA sequence FRT-*CmR*-FRT-B1006-J23116-*lacO*-B0032 before start codon of *ftsZ* in MG1655 | This study |
| MG-633Z | Inserting DNA sequence FRT-*CmR*-FRT-B1006-J23116-*lacO*-B0033 before start codon of *ftsZ* in MG1655 | This study |
| MG-634Z | Inserting DNA sequence FRT-*CmR*-FRT-B1006-J23116-*lacO*-B0034 before start codon of *ftsZ* in MG1655 | This study |

| | | |
|---|---|---|
| MG-635Z | Inserting DNA sequence FRT-*CmR*-FRT-B1006-J23116-*lacO*-B0035 before start codon of *ftsZ* in MG1655 | This study |
| MG-664Z | Inserting DNA sequence FRT-*CmR*-FRT-B1006-J23116-*lacO*-B0064 before start codon of *ftsZ* in MG1655 | This study |
| TC-29Z | Inserting DNA sequence FRT-*CmR*-FRT-B1006-ThrSen1.0-B0029 before start codon of *ftsZ* in TG | This study |
| TC-30Z | Inserting DNA sequence FRT-*CmR*-FRT-B1006-ThrSen1.0-B0030 before start codon of *ftsZ* in TG | This study |
| TC-31Z | Inserting DNA sequence FRT-*CmR*-FRT-B1006-ThrSen1.0-B0031 before start codon of *ftsZ* in TG | This study |
| TC-32Z | Inserting DNA sequence FRT-*CmR*-FRT-B1006-ThrSen1.0-B0032 before start codon of *ftsZ* in TG | This study |
| TC-33Z | Inserting DNA sequence FRT-*CmR*-FRT-B1006-ThrSen1.0-B0033 before start codon of *ftsZ* in TG | This study |
| TC-34Z | Inserting DNA sequence FRT-*CmR*-FRT-B1006-ThrSen1.0-B0034 before start codon of *ftsZ* in TG | This study |
| TC-35Z | Inserting DNA sequence FRT-*CmR*-FRT-B1006-ThrSen1.0-B0035 before start codon of *ftsZ* in TG | This study |

| | | |
|---|---|---|
| TC-64Z | Inserting DNA sequence FRT-*CmR*-FRT-B1006-ThrSen1.0-B0064 before start codon of *ftsZ* in TG | This study |
| TG-RFP | TG carrying pE-RFP | This study |
| TH-103Z-RFP | TH-103Z carrying pE-RFP | This study |
| TC-64Z-RFP | TC-64Z carrying pE-RFP | This study |

**Table S2. Plasmids used in this study.**

| **Plasmids** | **Description** | **Source** |
|---|---|---|
| pTKRed | $rep_{pSC101}$ $Spc^R$ lac-inducible expression; DNA repair protein from *E. coli*, RecA | Lab Stock |
| pCP20 | $rep_{pSC101}$ $Cm^R$ $Amp^R$ site-specific recombinase, FLP; temperature-sensitive variant of the phage λ repressor | Lab Stock |
| pE-RFP | ColE1 ori $Amp^R$ PJ23100 B0034-*rfp* | This study |

**Table S3. Primers used in this study.**

| **Primer name** | **Sequences (5’-3’)** |
|---|---|
| 109-lacO-0029-F | GCTAGCGGAATTGTGAGCGGATAACAATTCCTCTAGAGTTCACACAGGAAACC |
| 06-109-lacO-R | TCCGCTAGCACAGTCCCTAGGACTGAGCTAGCTGTAAAAAAAAAAACCCCGCCCTGTC |
| lacO-0029-Z-F | TCCTCTAGAGTTCACACAGGAAACCTACTAGATGTTTGAACCAATGGAACTTACCAAT |
| 109-lacO-RBS-R | ACTCTAGAGGAATTGTTATCCGCTCACAATTCCGCT |

| | |
|---|---|
| | AGCACAGTCCCTAG |
| YZ-Re-Z-F | CATCCCGTACACGGATCACG |
| YZ- Re-Z-R | CATAGTTACGCATCTGTGAGCGAT |
| TGDZ-UF | GAGTCTCAACGTCAGACACT |
| TGDZ-DR | GAGTCTCAACGTCAGACACT |
| 109-lacO-30-Z-F | GCGGAATTGTGAGCGGATAACAATTCCTCTAGAGA TTAAAGAGGAGAAATACTAGATGT |
| lacO-30-Z-F | TCCTCTAGAGATTAAAGAGGAGAAATACTAGATGT TTGAACCAATGGAACTTACCAAT |
| 109-lacO-31-Z-F | GCTAGCGGAATTGTGAGCGGATAACAATTCCTCTA GAGTCACACAGGAAACC |
| 109-lacO-32-F | GACTGTCTAGCGGAATTGTGAGCGGATAACAATTC CTCTAGAGTCACACAGG |
| lacO-32-Z-F | ATTCCTCTAGAGTCACACAGGAAAGTACTAGATGT TTGAACCAATGGAACTTACCAAT |
| 109-lacO-33-F | CTGTCTAGCGGAATTGTGAGCGGATAACAATTCCT CTAGAGTCACACAGG |
| lacO-33-Z-F | CAATTCCTCTAGAGTCACACAGGACTACTAGATGTT TGAACCAATGGAACTTACCAAT |
| 109-lacO-34-F | CTGTCTAGCGGAATTGTGAGCGGATAACAATTCCT CTAGAGAAAGAGGAGAAATACT |
| lacO-34-Z-F | AATTCCTCTAGAGAAAGAGGAGAAATACTAGATGT TTGAACCAATGGAACTTACCAAT |
| 109-lacO-35-F | CTAGCGGAATTGTGAGCGGATAACAATTCCTCTAG AGATTAAAGAGGAGAATACTAGA |
| lacO-35-Z-F | ATTCCTCTAGAGATTAAAGAGGAGAATACTAGATG TTTGAACCAATGGAACTTACCAAT |
| 109-lacO-64-F | GGACTGTCTAGCGGAATTGTGAGCGGATAACAATT CCTCTAGAGAAAGAGGGGAAAT |

| | |
|---|---|
| lacO-64-Z-F | AATTCCTCTAGAGAAAGAGGGGAAATACTAGATGT TTGAACCAATGGAACTTACCAAT |
| 06-116-lacO-R | CCGCTAGCATAGTCCCTAGGACTGAGCTAGCTGTC AAAAAAAAAACCCCGCCCTGTC |
| 116-lacO-R | CTCTAGAGGAATTGTTATCCGCTCACAATTCCGCTA GCATAGTCCCTAGG |

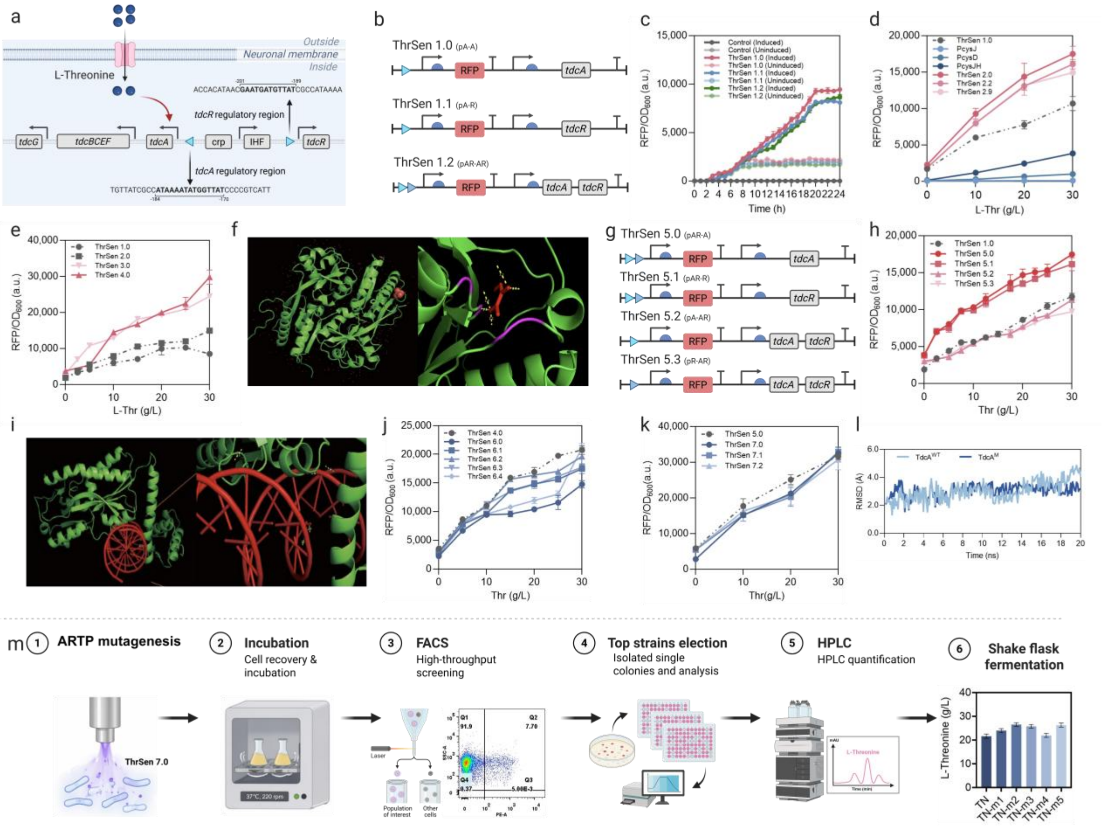


**Figure S1. Construction, optimization, and application of an L-threonine biosensor.**

a, Organization of the tdc operon and its regulatory mechanism. b, Design of the prototype L-threonine biosensor. c, Characterization of the L-threonine responsiveness of TdcA/TdcR and their cognate regulatory sequences. d, Screening of single-site variants within the TdcA ligand-binding domain. e, Characterization of combinatorial multisite variants. f, Structural analysis of a TdcA variant in complex with L-threonine. g, Strategy for reorganizing the TdcA/TdcR regulatory elements. h, Response profiles of the combinatorial biosensors. i, Schematic representation of TdcA DNA-binding-domain optimization. j, Characterization of DNA-binding-domain variants. k, Performance of the combinatorially optimized biosensor ThrSen 7.0. l, Structural stability analysis of the TdcA variants. m, Application of ThrSen 7.0 in combination with atmospheric and room-temperature plasma (ARTP) mutagenesis and fluorescence-activated cell sorting (FACS) to screen for L-threonine-overproducing strains.
